# Heritability estimation of subcortical volumes in a multi-ethnic multi-site cohort study

**DOI:** 10.1101/2024.01.11.575231

**Authors:** Christian Coffman, Eric Feczko, Bart Larsen, Brenden Tervo-Clemmens, Gregory Conan, Jacob T. Lundquist, Audrey Houghton, Lucille A. Moore, Kimberly Weldon, Rae McCollum, Anders J. Perrone, Begim Fayzullobekova, Thomas J. Madison, Eric Earl, Oscar Miranda Dominguez, Damien A. Fair, Saonli Basu

**Affiliations:** Division of Biostatistics, University of Minnesota, 100 Church Street SE, Minneapolis, 55455-0213, MN, USA; Department of Pediatrics, University of Minnesota, 100 Church Street SE, Minneapolis, 55455-0213, MN, USA; Department of Psychiatry & Behavioral Sciences, University of Minnesota, 100 Church Street SE, Minneapolis, 55455-0213, MN, USA; Department of Psychology, University of Minnesota, 100 Church Street SE, Minneapolis, 55455-0213, MN, USA; Masonic Institue for the Devloping Brain, University of Minnesota, 2025 East River Parkway, Minneapolis, 55414, MN, USA

**Keywords:** heritability, ABCD, SNP, fMRI

## Abstract

Heritability of regional subcortical brain volumes (rSBVs) describes the role of genetics in middle and inner brain development. rSBVs are highly heritable in adults but are not characterized well in adolescents. The Adolescent Brain Cognitive Development study (ABCD), taken over 22 US sites, provides data to characterize the heritability of subcortical structures in adolescence. In ABCD, site-specific effects co-occur with genetic effects which can bias heritability estimates. Existing methods adjusting for site effects require additional steps to adjust for site effects and can lead to inconsistent estimation. We propose a random-effect model-based method of moments approach that is a single step estimator and is a theoretically consistent estimator even when sites are imbalanced and performs well under simulations. We compare methods on rSBVs from ABCD. The proposed approach yielded heritability estimates similar to previous results derived from single-site studies. The cerebellum cortex and hippocampus were the most heritable regions (***>*50%**).

## 1 Introduction

Understanding the heritability of traits is essential for advancing early diagnosis and treatment across a spectrum of conditions. Genetic factors play a pivotal role in shaping an individual’s susceptibility to various traits, influencing outcomes ranging from physical characteristics to disease predispositions. Determining the extent to which certain traits are heritable can help untangle genetic and environmental contributions in order to help identify individuals at risk and tailor personalized interventions. Heritability has informed treatment approaches from traits amenable to medical intervention (such as cleft-pallette) to diseases requiring proactive healthcare strategies (such as Tay-Sachs). Understanding heritability is critical for developing targeted interventions, enabling early intervention, optimizing treatment efficacy, and potentially mitigating the impact of genetic predispositions on health outcomes. [1]

Endophenotypes, intermediate phenotypic markers between genetic variation and clinical manifestations, serve as crucial tools in mental health studies. These traits, more directly influenced by genetic factors than complex clinical outcomes, provide a mechanistic bridge, linking the genetic underpinnings of mental health disorders to observable behavioral and neurobiological characteristics. The utility of endophenotypes lies in their potential to dissect the complexity of psychiatric conditions into simpler, more tractable and heritable components for investigation. Compared to clinical manifestations, endophenotypes often exhibit larger effect sizes, are quantifiable, and manifest simpler genetic architectures. This makes them particularly valuable in discerning the genetic factors contributing to mental health outcomes, aiding in the identification of susceptibility genes and pathways.

Investigating the heritability of endophenotypes in adolescents holds particular significance as this developmental stage represents a critical period marked by the emergence of many mental health maladies [2, 3]. Many of these maladies are known to be heritable to varying degrees such as ADHD, major depressive disorder (MDD), and schizophrenia [4]. The genetic underpinnings of these maladies are complex and under investigation. Investigating their heritability during adolescence likely holds key genetic information to potentially inform targeted interventions and preventative strategies [5].

Regional subcortical brain volumes (rSBVs) are associated with psychiatric disorders, such as post-traumatic stress disorder (PTSD) [6] and major depressive disorder (MDD) [7] and are heritable [8]. Therefore, rSBVs can serve as valuable endophenotypes for mental health outcomes. Despite the pivotal role that rSBVs play as potential endophenotypes for psychiatric disorders, a notable gap exists in our understanding of their heritability in adolescents. While some studies have ventured into this domain, their limitations, such as small sample sizes or wide age ranges, have hindered comprehensive characterization. The intricacies of neurodevelopment during adolescence necessitate focused investigations to disentangle the unique genetic influences on rSBVs during this critical period. The existing body of research, though valuable, often lacks the precision needed to draw definitive conclusions about the heritability of rSBVs in adolescents[9]. Addressing this gap is imperative to gain a nuanced understanding of the genetic factors contributing to regional sub-cortical brain volumes during this crucial developmental phase, ultimately informing more effective interventions and diagnostic strategies for psychiatric disorders emerging in adolescence.

The Adolescent Brain Cognitive Development (ABCD) study is ideally suited to investigate the heritability of rSBVs in adolescents with its remarkably narrow age range. Additionally, ABCD has a remarkbly large sample size with over 10,000 subjects at the baseline age of 9-10 years [10]. The robust representation of diverse individuals in the study reflects the demographic richness of the United States, enhancing the generalizability of findings [11]. ABCD is uniquely designed to answer scientific questions addressed at the neurodevelopment of adolescents such as the heritability of rSBVs within at that age.

A challenge to estimating heritability in large multi-site studies like ABCD is that the study’s multi-site, multi-manufacturer design is confounded with the varying ethnic distributions across sites. The complexities of these site and batch effects must be addressed in order to estimate heritability effects [12]. Existing methods designed to adjust for these effects have resulted in inconsistent heritability estimates, as we will show in the methods section. Whereby inconsistent we mean the estimates fail to converge in a probabilistic sense to the true value for large sample sizes. These methods include Site Wise Demeaning (SWD), Adjusted Residual Adjustment, ComBat, CovBat, and meta-analysis approaches, each with its own unique strategy for addressing site effects but lacking considerations for genetic ancestry.

To overcome this challenge, we introduce a novel approach: AdjHE-RE (Adjusted Hasemann-Elston estimation with random site effects). This method offers consistent estimation of SNP-heritability by adjusting for site effects while controlling for the potential dependencies between site and race/ethnicity. Utilizing a method of moments approach, AdjHE-RE guarantees statistical consistency for heritability estimates [13].

This paper presents AdjHE-RE and describes tests of its reliability. We validate its consistency using synthetic data across various scenarios and apply it to the ABCD dataset, mirroring existing results in different populations. Our results reveal the enhanced consistency and reliability of AdjHE-RE across a wide range of scenarios, providing heritability estimates that align with established results. In the following sections, we delve into the development and validation of AdjHE-RE, offering a detailed account of its statistical framework and its practical implications in advancing our understanding of SNP-heritability in the context of regional subcortical brain volumes.

## 2 Methods

### 2.1 Genotype, Phenotype, and heritability

We assume a single quantitative phenotype (***Y*** ^∗^) following equation 1 with *n* observations of individuals from *K* subpopulations/ethnicties (*K* ∈ ℕ, with *n*_*k*_ denoting the number of observations in group *k* and 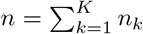, typed at *M* SNP positions. Assume a genotype matrix ***X***_*g*_ (*n* × *M*) where the *ij* th entry represents the number of allele copies at SNP position *j* for subject *i* and take the values 0, 1, or 2. The ***X***_*g*_ are observed from *K* clusters yielding ***X***_*gk*_ for each *k* ∈ {1, …, *K*} and are concatenated row-wise to make ***X***_*g*_. For each subpopulation (*k*) the allelic frequency at position *j* is given by *f*_*kj*_. The standardized genotypes are then given 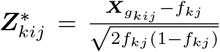. The genetic relatedness matrix (GRM, ***A***^∗^) is then defined as follows ***A*** ^∗^ =***Z*** ^∗^***Z***^′∗^*/M*. However, many analytical methods generally assume a homogeneous population and the GRM is estimated as follows where 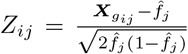, where 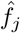 is the allelic frequency for the entire sample 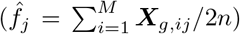. This means that in practice the following version of the GRM is generally estimated ***A***=***ZZ*** ^′^*/M*.

We define ***X***_*S*_ as an *n* × *p*_*s*_ matrix where the *ij* th entry is 1 if subject i was observed at site *j* and 0 otherwise codifying site membership. We define ***X***_*d*_(*n* × *p*_*d*_) as a set of covariates defining demographics such as sex and age and an intercept. The covariates can take on many forms such as categorical for sex and continuous for age. The columnwise concatenation of the demographic variables (***X***_*d*_) with the site membership matrix (***X***_*S*_) will be denoted as ***X***_*dS*_. Note that *p*_*d*_ *< p*_*s*_ *< n << M*.

A subset of *C* SNPs are assumed to causally affect the phenotype (***Y*** ^∗^) for all groups 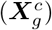. We assume a linear model where for the phenotype of the ith subject

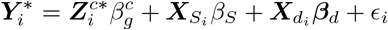

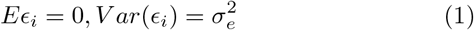

In order to quantify the genetic contribution, we assume 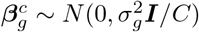. We define 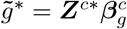 [14], then Eqn 1 reduces to

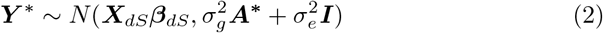

Reexpressing the model in terms of random genetic effects we have (assuming the covariance between the genetic effects and unmodeled error (*ϵ*) is 0 and that the genetic variance coefficient 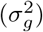 is constant across all *K* populations. The heritability *h*^2^ of the trait ***Y***^∗^ is defined as 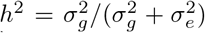. We are interested in estimating the parameters 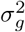 and 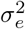. In the following methods, we achieve this estimation through residualizing the phenotype either on some of the covaraites affecting the mean 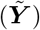 where the estimated coefficients are also denoted with a tilde. Other methods residualize the phenotype on all of the covariates affecting the mean simultaneously (***Y***). Lastly, we refer to the phenotype in capitalized form when it is a random variable, but in lower case when it is observed (such as when writing out the empirical estimators).

#### 2.1.1 Heritability Estimation Approaches

In general, we do not know which variants are causal. For heritability estimation, we consider ***A*** instead of ***A***^∗^. Without loss of generality, we assume that there are no site effect present. In absence of site effects we have ***Y***^∗^ ∼ 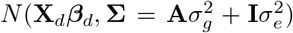. For Restricted Maximum Likelihood Estimation (REML) we consider the log-likelihood of the data that is given by:

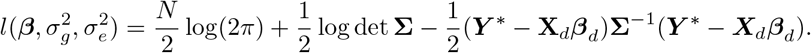

The estimation of the mixed effect model mentioned above is performed through maximum likelihood estimation. The software GCTA [14] uses the iterative restricted maximum likelihood (REML) algorithm to estimate the variance components 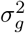 and 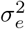 in the model described above and gives an estimate of heritability by 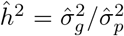, where 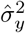 is the estimated phenotypic total variance 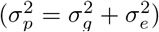.

However, this REML-based estimation for a mixed-effect model can easily get computationally intractable as the sample size increases. Recently, there have been attempts to generate computationally scalable algorithms to implement mixed models on large scale data [15–17]. However, these approaches still encounter computational challenges on large biobanks. Moreover, even subtle population substructure can significantly impact the heritability estimation [18].

The method of moments (Haseman-Elston regression [19], LDscore regression [20], MMHE [21]) approaches are another set of widely used methods for estimating heritability *h*^2^ under Equation 1. We will next provide short overview of the Hasemann-Elston method as it is most similar to the newly proposed estimator.

In Haseman-Elston (HE) Regression, we generally assume that the GRM is normalized with its diagonal entries all equal 1 and **Y** is centered and that Equation 2 holds. One of the classical moment estimators for *h* ^2^ comes from the least squares regression coefficient for regressing **Y** _*i*_ **Y**_*i′*_ on **A**_*ii′*_ for all *i < i* ^′^. This is because Equation 1 implies that *E*(**Y** _*i*_**Y**_*i′*_ |**A**) =*h* ^2^**A**_*ii′*_ for *i* ≠ *i* ^′^. The heritability can be estimated from the following equation:

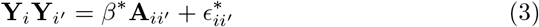

Note *β* ^∗^ = *h*^2^ is the heritability parameter. The corresponding estimator for *h*^2^ is 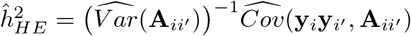.Note that

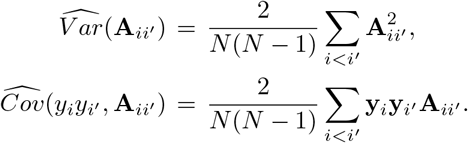

[22] used least squares in this way to estimate 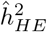. This approach is also referred to as Haseman-Elston (HE) regression [19].

The above approaches generally assume a homogeneous population. In a sample of diverse ancestry or even in presence of subtle substructure, these existing methods could produce very biased estimates of heritability. The standard approach is to adjust for principal components (***P Cs***) of the GRM **A** as covariates in Equation 1 and perform REML estimation[14]. Lin et al [23] proposed AdjHE approach that proposes a product PC approach to correct for the population stratification effects in both mean and variance of **y**.

Under AdjHE model if we assume the *K* subpopulations can be completely separated by *K* PCs then 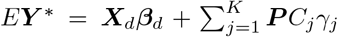 and 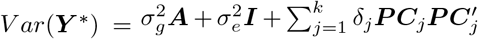 . Letting *K* be the number of genetic principal components being adjusted for, let 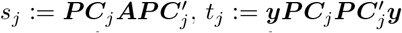, and 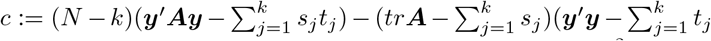, the AdjHE method of moments solution proposed the following 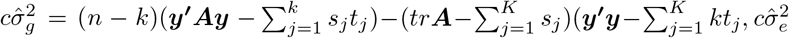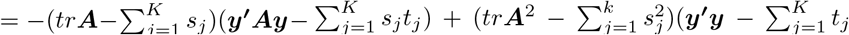. The heritability is then estimated as 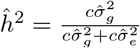. [23]

The adjHE approach assumes that fixed covariates and the genetic random effects are uncorrelated. However, when we expect site effects, they are likely to be correlated such as with the ABCD study because of the differential ethnic distribution across sites.

### 2.2 Existing methods controlling for site effects in neuroimaging studies

There are three ways to control for site effects: a) The first is to perform a stepwise regression in which the first step involves residualizing the phenotype against site membership followed by AdjHE estimation on the residualzied phenotypes; b) The second method involves adjusting for the site effect in one or both of the two regression steps that are inherently part of AdjHE estimation c) The third method involves making separate estimates for each site and pooling them in a meta analysis. The previously existing methods generally involve regressing out the site effects in a prior step. To compare how each method affects SNP heritability estimation, we first transform the phenotype as dictated by the given method and then estimate heritability using AdjHE which yields the heritability estimate. The heritability estimate is given by 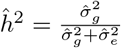. Note that there are additional methods used to address site effects using preprocessing procedures [24, 25] and nonlinear transformations[12] that aren’t considered here.

Using method of moments under the umbrella of M-Estimation (Generalized estimating equations) theory we will evaluate consistency by looking at the defining equations, evaluating the expectation of the equation set to 0 (to determine the true parameter) and determine whether it is the parameter of interest. In this context, the conditions for consistent estimation are met when the genetic effects are unaltered when residualizing out the site and demographic effects on the mean.

#### 2.2.1 Method a1: Stepwise Adjustment Approach: Site-Wise De-meaning (SWD)

SWD adjustement and heritability estimation proceeds in 5 steps: 1) Regressing out the mean effects of site membership, 2) regressing out demographic effects on the mean 3) regression on the variance components and plug-in estimation of *h*^2^ [26], 4) estimating the variance components attributable to the GRM and Identity matrices 5) plug-in estimation for heritability. We represent this process of estimation through the use of estimating equations. The first moment estimating equation 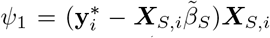. The second estimating equation for the remainder of the mean (excluding the SNP effects) is 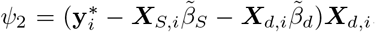. The third estimating equation establishes the residualized phenotype 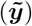 as 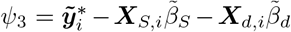. The fourth estimates the coefficients corresponding to the variances attributable to the GRM and identity matrices 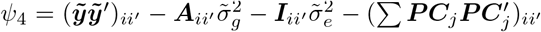. The estimated heritability is given by the final estimating equation 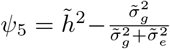.

Looking at the expectation of residualizing in 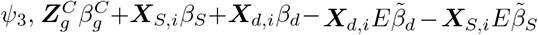. When 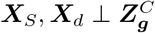 the expectation simplifies to 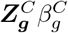 and means that 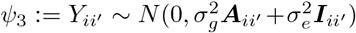. It would follow that an OLS estimate on the 2nd moment using the GRM and the Identity matrix would yield consistent estimates of 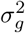 and 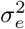 by the properties of OLS estimators. Lastly, by the continuous mapping theorem, the estimate of *h*^2^ is consistent.

#### 2.2.2 Method a2: COMBAT/COVBAT

Combat utilizes an empirical Bayes estimator to adjust for site effects [27, 28]. Heritability estimation follows in multiple steps: 1) estimate and remove the site specific phenotype mean and variance as the posterior conditional means 2) estimation and subtraction of demographic effects 3) OLS estimation of variance components along with plug-in estimation of *h* ^2^. Since empirical Bayes estimators are consistent, we know that the projection converges in probability to the frequentist estimate of site specific mean. Similarly, since we are assuming that the measurements have equal variance across sites, the COMBAT estimate of the site variances will converge in probability to the identity matrix scaled by 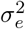. In other words, COMBAT adjustment is asymptotically equivalent to the SWD method unless the other effects (***X***_*d*_*β*_*d*_) are controlled for in the estimation of the site effects in which case it is asymptotically equivalent to Adjusted residual adjustment, which is discussed in the next section.

Covbat is an extension of Combat that additionally adjusts for the covariance of the site effects amongst multivariate traits. Covbat adjustments are done after first performing Combat and would therefore suffer from the same inconsistency of Combat. Covbat estimates the covariance of the scanner effects from a PCA decomposition of the Combat residualized phenotypes. For heritable traits, there is additional covariance from the GRM that the PCs would capture and thus adversely affect the estimated heritability.

#### 2.2.3 Method b: Adjusted residual adjustment

Adjusted residual adjustment and estimation proceeds in four steps: 1) residualize the phenotype on the site and additional covariates simultaneously 2) OLS estimate of variance components on residuals with plug-in estimation for *h*^2^. 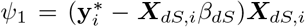. The second estimating equation establishes the residualized phenotype (**y**) as 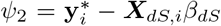. The third estimating equation estimates to coefficients corresponding to the variances attributable to ***A*** and ***I*** 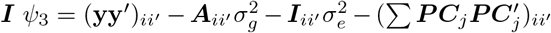. The estimate of heritability is given by the final estimating equation 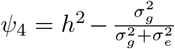. Looking at the expectation of residualizing in 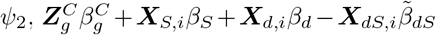 . This leaves the genetic contributions in tact when ***X***_*dS*_ ⊥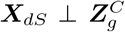. Under these conditions, after residualizing 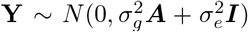 and the following OLS estimate from *ψ*_4_ and plug-in estimate for *h*^2^ in *ψ*_5_ are consistent estimators given the 2nd moment is correctly specified by properties of OLS estimation.

#### 2.2.4 Method c: Meta analysis

Meta-analysis approaches are considered where the general procedure involves heritability estimation for each site followed by a weighted average of the site-specific estimates. For fixed effects meta-analysis approaches, the inverse variance weighted mean is known to minimize the variance of the weighted averages and has been selected here for comparison [25, 29]. This is commonly chosen for meta-analyses in which the only error in the estimate is due to the sampling variance of each study [30]. AdjHE and GCTA estimates are unbiased for single-site studies [23]. Using M-estimator theory then we have 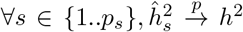 . Therefore by the continuous mapping theorem, 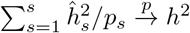 making it a consistent estimator for both AdjHE and GCTA as long as the model for each of the separate site estimates are properly defined.

## 3 New Method: Modeling site as a random effect

The demographic variables (***X***_*d*_, sex, and age), except estimated genetic ancestry (***PC***), are reasonably assumed to be linearly independent of site membership (***X***_*S*_). We apply *Q*_*d*_ (*Q*_*d*_ projection not including PCs) such that the first moment *EQ*_*d*_***Y*** = 0 assuming *Eβ*_*G*_ = *Eβ*_*S*_ = *Eϵ* = 0. The second moment of the transformed phenotype is given by 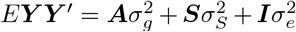 (assuming the independence of *β*_*G*_, *β*_*S*_, *ϵ*), where *m* is the number of SNP’s. Estimates are derived from an OLS estimator on the 2nd moment (Full details are included in Appendix A) yielding the solution

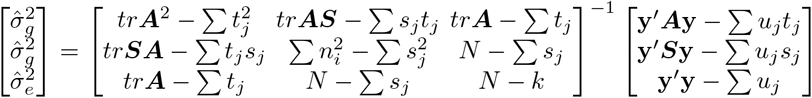

where 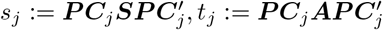, and 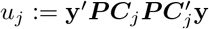

Assuming that the model (Equation 1) is correctly specified, specifically that the first and second moments are accurately specified, the estimates of the variance parameters obtained using M-estimation theory are consistent [13]. Assuming the additional assumptions used to get the simplified solution above, such as the demographic variables, but not necessarily the PCs, are orthogonal to the site membership (**X**_*S*_) or when there is not a fixed effect contribution from the PCs. Following the delta method in Appendix D it is evident that the variance of the heritability is identical to that of the fixed effects model [21, 23].

## 4 Results

We assessed each of the estimators over a wide array of simulation setups (Sections 4.1, 4.2, 4.3, and 4.4). Then, we compare results on ABCD-derived subcortical volumetric data (Section 4.6).

### 4.1 Simulations

A representation of the Monte Carlo simulation procedure following the Balding-Nichols method [31] is shown in Figure 1. Each simulation was replicated 100 times. Each simulation had the following set of variables in common (unless otherwise noted): # SNPs = 20000, *θ*_*alleles*_ = 0.1, proportional causal SNPs = 0.02, no interaction between SNP effect and site, 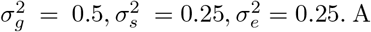. A single continuous covariate was simulated to represent demographic variables that might act as a fixed effect on the trait such as age or sex. Simulations had either 2 or 25 sites, 1 or 2 genetic clusters, and equal or heterogeneous distributions of ancestries across sites.

**Fig. 1.**
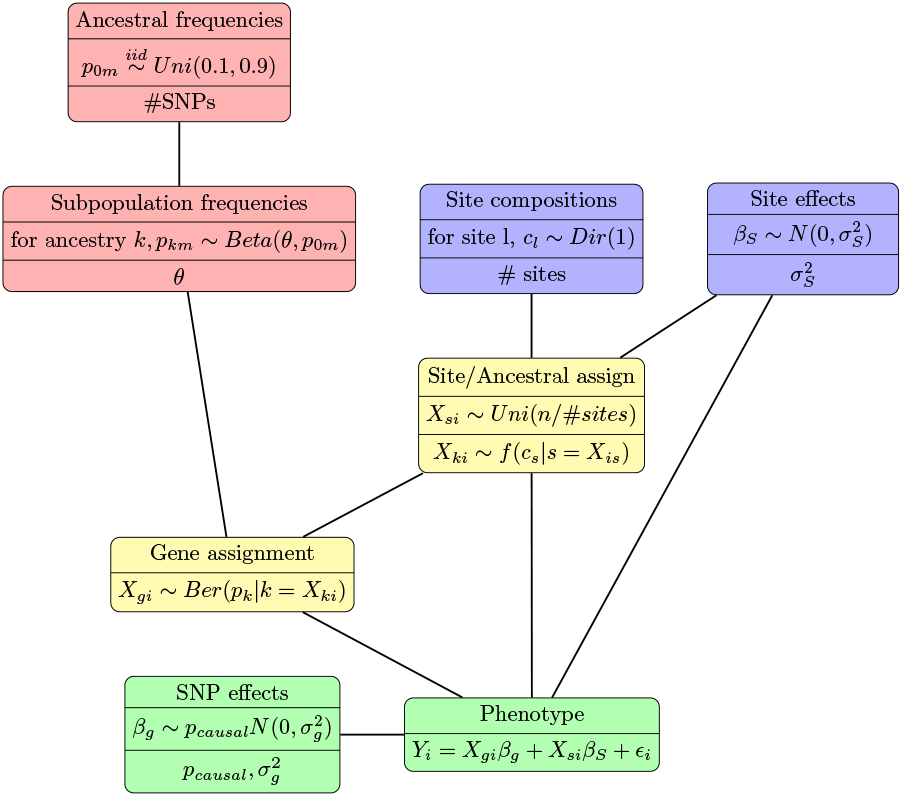
Diagram of the method of simulating phenotypes from multiple sites that depend on genotypes that may be clustered.

We simulated allele (*m*) frequencies (*p*_0*m*_) for ancestors common to all subjects iid from a *Uniform*(0.1, 0.9) yielding a vector of allele frequencies,*p*_0_. Then, the allele frequencies for each genetic cluster (*k*) was drawn from a *Beta*(*p*_0*m*_(1 − *θ*)*/θ*, (1 − *p*_0*m*_)1 − *θ*)*/θ*) distribution with *θ* = 0.1, yielding allelic frequencies for each genetic cluster (*p*_*k*_). (See red blocks in Figure 1)

The composition of each site was either forced to contain equal numbers from each cluster or sampled from an equiprobability Dirichlet distribution (*Dir*(1)). Site effects were drawn from a standard normal or set as fixed effects (Figure F3) and scaled by 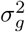. (See blue blocks in Figure 1). Subjects were assigned to a genetic cluster based on the already sampled site composition. Conditional on the subject’s cluster, the genotype was sampled from a *Binomial*(2, *p*_*k*_) (*f*_*g*_ in the diagram). (See yellow blocks in Figure 1).

SNP effects were simulated from a standard normal scaled by 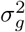 conditional on being selected as a causal SNP which followed independent Bernoulli(*p*_*causal*_) distribution at each position. The phenotype was then calculated using the data-generating mechanism. (See green boxes in Figure 1). For each simulated dataset, we conducted each estimating procedure. Multiple variations of this simulation setup were run varying how the population clusters were distributed across sites and how the site effects were distributed as is discussed in Sections 4.2, 4.3.

The results of each simulation experiment are reported in the following sections. Overall, the proposed estimator showed unbiased estimates for simulations of multiple genetic clusters unevenly sampled over 2-25 sites including in the presence of site heteroskedasticity, becoming less biased with a larger number of sites (Figure 2. The method is also unbiased under simpler conditions (Figures F1) except for simulations consisting of persistent, evenly spaced fixed site effects under which all estimators showed some level of bias (Figure F3). The naive meta-analysis estimators (“nAdjHE”, “nGCTA”) showed similarly unbiased results. In general, the simulations agreed with theoretical considerations in section 2.2 that suggested negating the site effects without considerations for population imbalance or the GRM would result in increasingly biased heritability estimates. Given that the bias of these estimators increases with the imbalance between sites. Both GCTA and AdjHE showed unbiased estimates and similar variance for a homogeneous population at a single site (Figure F1). Most estimators provided nearly unbiased estimates for balanced sites with homoskedastic error with Combat and Covbat adjustments showing an appreciable negative bias (Figure F2).

**Fig. 2.**
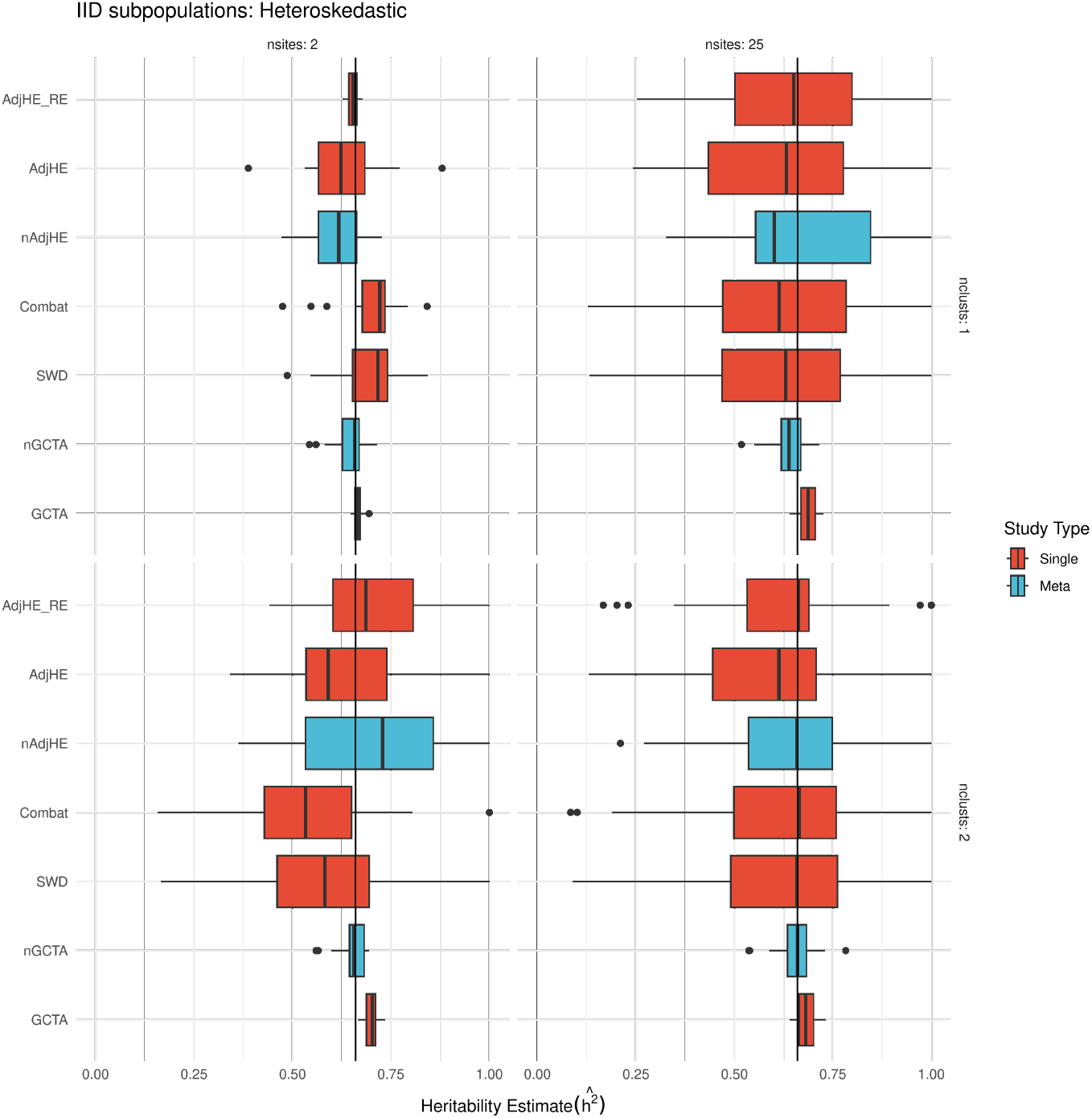
Estimated heritability with an increasing number of sites (columns) and clusters (rows) where *h*^2^ is fixed at 66%. The total phenotypic variance was simulated 50% from genetic data, 25% from sites, and 25% from noise making the simulated heritability 66% (vertical black line).

### 4.2 Simulation setup 1, Homoskedastic site effects

Site were simulated to contain a perfectly balanced number of subjects from each cluster. The first trivial setup involved the estimation of a homogeneous population from a single site. Since it only involved one site it lent it restricted estimation to GCTA or AdjHE methods. These simulations were repeated over an array of 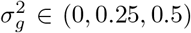 which are represented as faceted column plots in Figure F1. The true simulated heritability in this figure is represented as a horizontal black line and the distribution of estimation results from each procedure are represented as a boxplot with estimations following outside 1.5 multiples of the interquartile range represented as points. Both AdjHE and GCTA have medians that are almost indistinguishable from the true heritability and with similar distributions about the simulated heritability under all combinations of variance contributions from site, genetics, and error.

Homoskedastic simulations were continued for 2 and 25 sites (columns) as well as 1 and 2 genetic clusters (rows) (Figure F2). Estimates in these plots are colored according to whether they are emulating a single study with multiple sites (red) or a meta-analysis (blue) where each is comprised of single-site studies. Most of the estimates are close to the simulated heritability except when 25 sites and 2 clusters were simulated when the ad hoc adjustments represented by Combat and SWD significantly underestimate the simulated heritability. The variances remain consistent across the number of sites and the number of population clusters with GCTA-related methods (GCTA and nGCTA) maintaining the smallest.

Lastly, simulations with imbalanced sampling Figure unequal sampling from the population clusters are visualized in Figure 2. The clusters in this instance were IID from a discrete uniform distribution so they are not generally equally distributed across sites. Under this set of conditions, most estimates were unbiased, the ad hoc methods (Combat and SWD) had the largest negative bias, especially for a large number of sites.

### 4.3 Simulation 2 Heteroskedastic site variance

The next set of simulations involved keeping the same setup as discussed in Section 4.2 except with the addition that site variances (*δ*_*s*_) were sampled iid from *Gamma*(4,4). The results of these simulations can be see in Figure 2. In addition, the variances tended to increase with the number of sites and the number of population clusters with GCTA-related methods maintaining the smallest variance.

### 4.4 Simulation 3, Fixed site effects

The same simulation setup as from section 4.2 was repeated, but where the site effects were uniformly spaced about 0 and kept as a fixed effect across all replicates scaled to 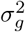. The results of this simulation setup are shown in Figure F3. Under the assumption of fixed site effects, both AdjHE estimators showed significant positive bias, and the GCTA meta-analysis (nGCTA) significant negative bias. None of the estimators under this condition were unbiased, but the ad hoc adjustments and GCTA estimator showed the least bias. The meta-analysis estimators (nGCTA, nAdjHE) both showed larger variances than the other estimators.

### 4.5 ABCD Data and Processing

The ABCD study is comprised of N=11,572 9-10 year-olds. ABCD is (parentheses describe the range across sites) 48% female (range 43-61%), 52% White (range 8-87%), 15% Black (range 0-54%), 2% Asian (range 0-13%), 20% Hispanic (range 2.3-74%), and 11% Other (range 3.6-23%) (See Table B1). Since the study is a secondary analysis of a data repository, IRB approval was not required. Instead, oversight is governed jointly by the University of Minnesota, the NIMH data archive, and the ABCD consortium. A data use agreement for the ABCD was authorized by UMN, signed by the investigators, and approved by the NDA after review.

#### 4.5.1 Genomic Data

All subjects were assayed using the Smokescreen™ Genotyping array [32] which includes more than 300,000 SNPs. Genotypes were phased and imputed on the Michigan imputation server https://imputationserver.sph.umich.edu/index.html#! [33]. Automated imputation QC steps were utilized on the Michigan server and additional QC steps were implemented on the imputed data utilizing Plink (v1.90b6.24, [34]). such as filtering the minor allele frequency to 5%, heterozygosity= 3SDs, selecting only single base pair replacement mutations, Hardy-Weinberg equilibrium = 1*e* −6, and a sliding window (200bp window size with 50 bp steps) independence of 0.25, imputation *R* ^2^ *>*0.99 . GRMs and eigendecompositions were then computed using Plink. The full dataset was filtered to remove both explicit and hidden family structures. After QC’ing the final sample size was N=7,234 spread over 20 sites and number of SNPs was 1,292,075.

#### 4.5.2 Image Acquisition, Preprocessing, and parcelation

Data acquisition is fully described at on the (https://abcdstudy.org/images/Protocol_Imaging_Sequences.pdf) [35]. fMRI data were accessed through the fast track portal 2.0 release. T1-weighted 3D MP-RAGE sequence with 1mm isotropic spatial resolution, TR/TE/TI = 2500/2.88/1060, and 8°flip angle. Parcelations were done following the DCAN ABCD-BIDS pipeline [36]. Anatomical data are registered to an MNI template, masked, denoised, and corrected for bias. Segmentations are then generated through Freesurfer [37]. Data is registered first to CIFTI template then to MNI followed by reporting of parcellated volumes [36].

### 4.6 Real data (ABCD)

#### 4.6.1 Heritability Estimates

Estimators were compared on subcortical volumes derived from the ABCD dataset. Each estimation controlled for a fixed effect from age and sex. The number of PCs was estimated to be 20 for the full dataset based on the stabilization of heritability estimates on global head volume as a function of the number of PCs for the global brain volume and a convergence of most of the estimates. Analysis was carried out on data from the single largest site (Figure G4), on reported European ancestry across all sites (Figure G5), and for the entire dataset (Figure G6).

Estimates carried out for subjects of European ancestry across all sites (n= 3669) across all phenotypes are shown in Figure G5. European heritage was chosen as it provided the largest sample size this yielded an average of 280 observations at each site with a minimum of 16 (See Table B1). Since this subset reduced the number of reported ancestries present in the data, the estimates were carried out with 0, 10, and 15 PCs. The estimates were relatively stable across the number of PCs for all estimators, especially between 10 and 15 PCs. In general, the meta-GCTA (nGCTA) method showed the largest heritability with GCTA estimating the least and all other estimators falling somewhere in between at roughly 10% heritability.

Estimates were carried out for site 16 (n= 622) across all reported ancestries and phenotypes. Site 16 was chosen because it had the largest sample size, note that the mean sample size is ≈329 with a standard deviation of 162. At Site 16 the distribution of races was 81% White, 12% Hispanic, 5.8% Other, 1.1% Black, and 0.2% Asian subjects (See Table B1). Since these estimates were carried out on a subset of the population, estimates were compared across a range of PCs (0-30). These estimates are visualized in Figure G4. Since these estimates were based on a single site only GCTA and AdjHE estimators were used. GCTA estimates are uniformly smaller than the AdjHE estimates and as the number of PCs are increased a general decrease in the number of large heritability is reported. The highest heritability estimated was for the Cerrebellum cortices.

Estimates for the full dataset (n= 7,234) across all phenotypes are shown on the brain in Figure 3 and as a scatter plot in Figure G6. The AdjHE methods estimate a larger heritability and the meta and ad hoc methods fall in between. Most estimates fell below 20% heritability with some (Hippocampus and Cerebellum cortex) being near or above 50%.

**Fig. 3.**
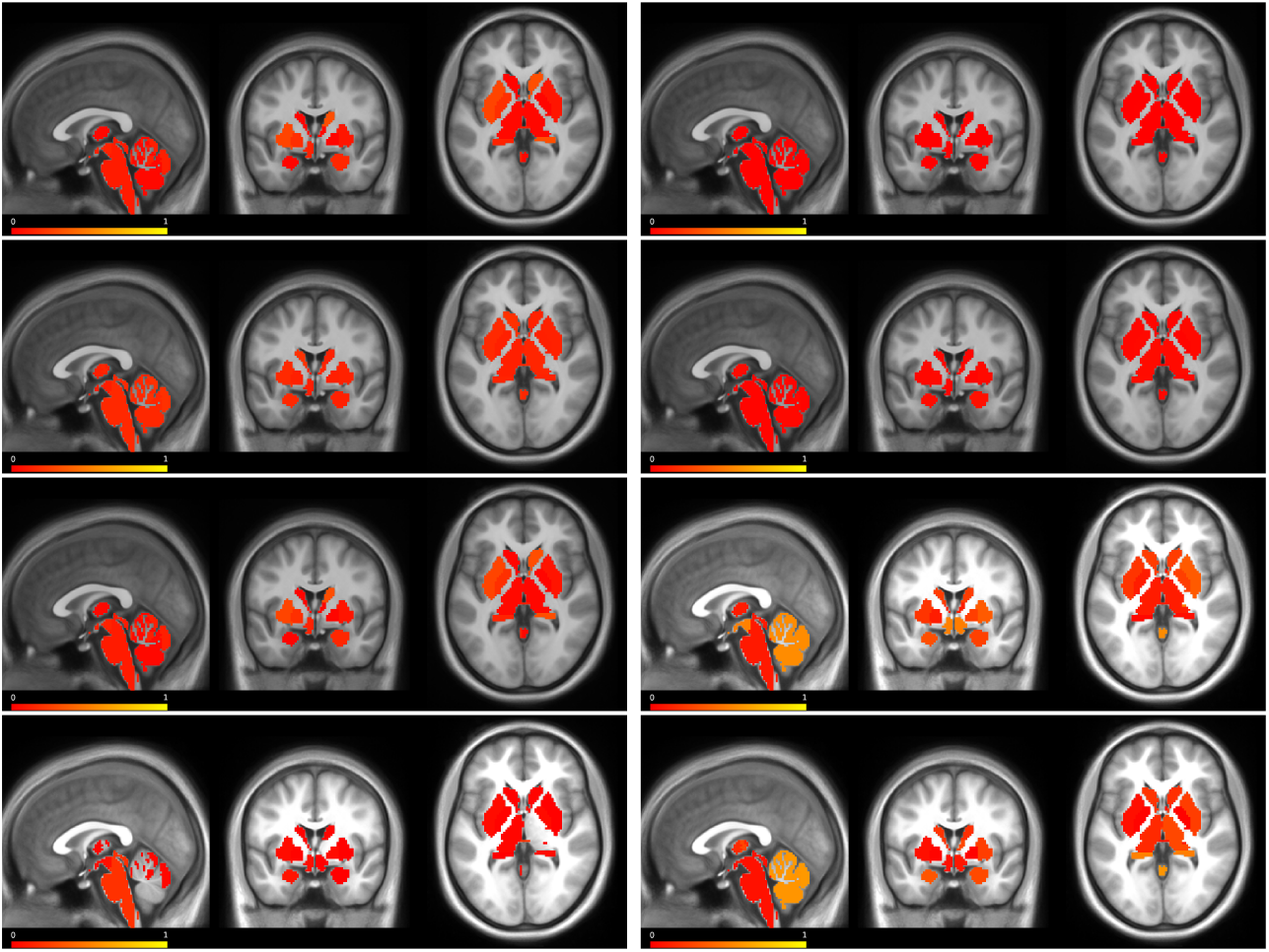
Heritability on the subcortical volumes from the full ABCD dataset visualized on the Conte T1 averaged brain. The heritability is reported as a heat map with red representing low heritability and yellow representing high heritability overlain on central axial, coronal, and sagittal slices, respectively. From left to right, top to bottom images represent estimates from AdjHE with fixed site effects, GCTA with fixed site effects, AdjHE meta-analysis, GCTA meta-analysis, SWD, Combat, Covbat, and AdjHE estimating site as a random effect.

### 4.7 Estimation efficiency

Comparisons of analysis time are compared in Table G2 and visualized in Figure 4. The complexity of GCTA estimation, as well as both AdjHE estimators, is *O*(*N* ^3^). For GCTA the complexity is based on the required GRM matrix inversion. For the AdjHE methods, the most complex operations are solving the matrix multiplication steps for solving the second moment equation. However, from profiling (Table G2) it is apparent that the AdjHE random variable estimator remains a consistent 3-4x faster than the GCTA algorithm and the AdjHE fixed effect model remains approximately 20x faster than the GCTA method. While the AdjHE methods provide a speed up it is at the cost of utilizing more RAM, especially with an increasing number of principal components since the outer product of the PC loading vectors is included for each PC modeled.

**Fig. 4.**
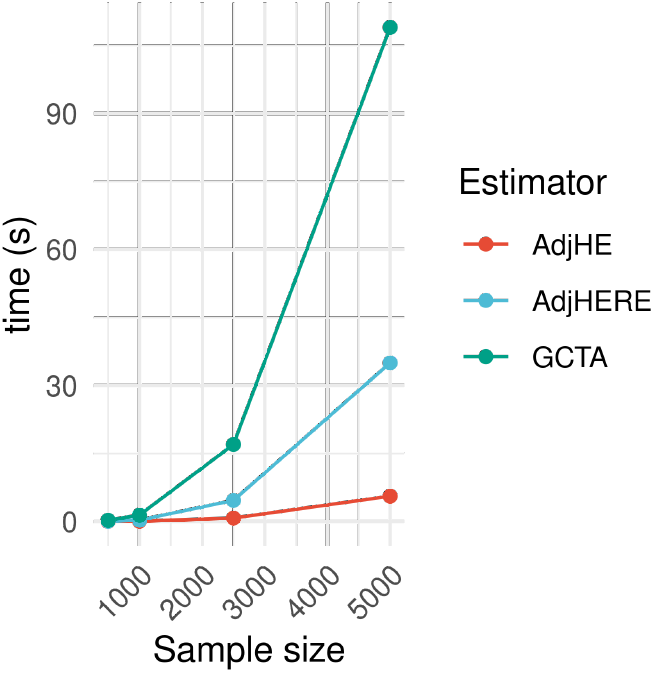
Computational wall time as a function of increasing sample sizes. (Red) computational time for AdjHE estimation, (Green) computational time for AdjHE with random site effects estimation, (Blue) computational time for GCTA estimation.

## 5 Discussion

Overall, this study illustrates that AdjHE-RE can yield unbiased estimates of SNP heritability for balanced and unbalanced data sets arising from multi-site studies, including in the presence of genetic clustering (such as with multiple ancestries). Additionally, we identified multiple rSBVs with significant heritability in adolescents, including a fair number that are associated with psychological disorders validating their role as endophenotypes. Along with this, the noted computational efficiency makes phenome-wide heritability analysis feasible on large datasets as are common in neuroimaging studies (Figure 4). AdjHE-RE as well as a wrapper for all of the other methods compared here are available publicly via the MASH toolbox on Github. The toolbox offers a command line tool for batch scripting as well as a Python interface for interactive computations.

Our work has resulted in two contributions to imaging genetics. First, we presented an extension to AdjHE estimation to adjust for site effects in heritability estimation which yields statistically consistent estimates. Theoretical and simulated comparisons corroborated the usefulness of the new estimator over a wide domain of simulated possibilities, including differing numbers of sites and genetic clusters, though comparisons were limited strictly to linear transformations (Section 4.1). AdjHE-RE was consistent even under non uniform error with imbalanced sites for 1-2 population clusters and 2-25 sites (Figure 2). The variance of AdjHE-RE was noted to decrease with the number of sites as is consistent with a more precise estimate of the variance of the random site effect. In addition, AdjHE-RE is up to four times faster than REML estimation (Figure 4). Overall, these features make AdjHE-RE amenable to effectively estimating heritability on multi-site studies such as ABCD.

AdjHE-RE has methodological limitations. The assumption that site effects behave as random effects is a simplification of the roll scanner effects and procedures might affect a measurement, and while it has proven effective in mitigating confounding factors, it may not fully capture all nuances in the data. Additionally, the current implementation lacks the capability to estimate nested confounding factors, such as scanner effects, which may introduce additional variability in the heritability estimates. Recognizing these limitations, we acknowledge that our method represents a step forward, but further refinement and exploration are necessary to address more intricate sources of confounding in multi-site studies.

Second, we estimated heritability using AdjHE-RE for rSBVs in ABCD and found multiple highly heritable regions, some of which are associated with psychological disorders (Figure 3). For example, the cerebellum had an estimated heritability of around 60% and is associated with anxiety [6]. The hippocampus with an estimated 50% heritability is associated with anxiety, major depressive disorder, and schizophrenia [7, 38]. These estimates share similarities with previous findings from PING (n = 500, age range = 3-21) and UKBiobank (n = 9000, age range = 40-69) despite very the large differences in sample demographics (mainly age and ethnicities) [9]. The largest estimates reported here, which were 60% for the cerebellum cortex and 50% for the hippocampus, largely agree with UKBiobank SNP-heritability estimates (63% and 50%, respectively) [9]. Additionally, the cerebellum has a major role in motor function and is thus understood to scale with height, a trait that has a well-established heritability above 60% [39]. The estimated heritability for amygdala volume is 35% compared to UKBiobank which was 23-42% [9]. GWAS estimates on the UKBiobank set a range of 20-60% heritability for T1 and T2 weighted subcortical structures [40]. Some weakly heritable traits were much lower than in previous reports. For example, estimates on the brain stem were 10% as compared to 82% from UKBiobank studies [9]. Differences might arise from age differences given UKBiobank is comprised of adults aged 40-69 years old compared to the ABCD sample which is 9-10 year-olds. The dramatic differences in brain stem estimates is additionally affected by “mid-brain shrinkage” that would be observed in UKBiobank and not in ABCD [41, 42].

One notable limitation in our study lies in the demographic constraints of the Adolescent Brain Cognitive Development (ABCD) dataset, which focuses exclusively on 9-10 year-olds residing near facilities with the requisite capabilities for inclusion in the study. This geographical and age restriction introduces a potential sampling bias, limiting the generalizability of our heritability conclusions to this specific cohort. The overrepresentation of individuals residing close to research facilities might introduce biases that do not capture the full diversity of the population, particularly underrepresenting rural communities. Consequently, caution is warranted when extrapolating our findings to broader demographics.

Due to an increasing number of multisite studies, being able to control for site effects will grow increasingly relevant. We plan to increase the capabilities of AdjHE. We plan to extend this method into multivariate modes of analysis which have also shown promise in neuroimaging studies [43]. We also plan to create a simulation toolbox to standardize testing of new methods for imaging genetics method controlling for site effects. For supplementary information please see the appendix.

## Supplementary information

For supplementary information please see the appendix.

## Additional information

### Data availability

All data is available at https://abcdstudy.org/.

### Code availability

The estimation tools are available at https://github.com/coffm049/Basu_herit. Genetics pipeline tools are available at https://github.com/coffm049/ABCD_geno_QC.

### Authors’ contributions

C.C. and S.B. designed the research; C.C. contributed all tools analysis and genetics processing; E.F. processed imaging data; C.C. wrote the manuscript; all authors aided in preparation and revision of manuscript.

### Ethics approval

Because the study is a secondary analysis of a data repository, IRB approval was not required. Instead, oversight is governed jointly by the University of Minnesota, the NIMH data archive, and the ABCD consortium. A data use agreement for the ABCD was authorized by UMN, signed by the investigators, and approved by the NDA after review.

### Consent to participate

Not applicable

