## Supplemental file for "Heritability estimation of subcortical volumes in a multi-ethnic multi-site cohort study"

### Appendix A Full derivation of AdjHE with random site effect

Estimates for this model will be derived from minimizing the 2-norm on the 2nd moment equation.

$$(\hat{\sigma}_g^2, \hat{\sigma}_g^2, \hat{\sigma}_e^2, \hat{\delta}_1, ..., \hat{\delta}_k) = argmin_{\delta_j,\sigma} yy' - (\begin{bmatrix} A & S & I & P_1P_1' & \dots & P_kP_k' \end{bmatrix} \begin{bmatrix} \sigma_g^2 \\ \sigma_g^2 \\ \sigma_e^2 \\ \delta_1 \\ \vdots \\ \delta_k \end{bmatrix}$$

Evaluating the 2-norm

$$Norm = yy' - (\sigma_g^2 A + \sigma_g^2 S + \sigma_e^2 I + \sum \delta_j P_j P_j')$$

$$\text{Let} G := \sigma_g^2 A + \sigma_g^2 S + \sigma_e^2 I + \sum \delta_j P_j P_j'$$

$$\nabla = (\frac{\delta}{\delta\sigma_g^2}, \frac{\delta}{\delta\sigma_g^2}, \frac{\delta}{\delta\sigma_e^2}, \frac{\delta}{\delta\delta_1^2}, ..., \frac{\delta}{\delta\delta_k^2})$$

$$\nabla Norm = (\frac{\delta Norm}{\delta\sigma_g^2}, \frac{\delta Norm}{\delta\sigma_g^2}, \frac{\delta Norm}{\delta\sigma_e^2}, \frac{\delta Norm}{\delta\delta_1^2}, ..., \frac{\delta Norm}{\delta\delta_k^2})$$

$$G'G = (\sigma_g^2 A + \sigma_g^2 S + \sigma_e^2 I + \sum \delta_j P_j P_j')'(\sigma_g^2 A + \sigma_g^2 S + \sigma_e^2 I + \sum \delta_j P_j$$

Setting the derivative of the Norm to 0

$$\nabla Norm = \nabla GG' - 2yy'\nabla G = 0$$

$$\nabla GG' = 2yy'\nabla G$$

Arranging this into matrix form

$$\nabla GG' = \begin{bmatrix} 2\sigma_g^2 trA^2 + 2A(\sigma_g^2 S + \sigma_e^2 I + \sum \delta_j P_j P_j') \\ 2\sigma_g^2 trS^2 + 2S(\sigma_g^2 A + \sigma_e^2 I + \sum \delta_j P_j P_j') \\ 2\sigma_e^2 trI^2 + 2I(\sigma_g^2 S + \sigma_g^2 A + \sum \delta_j P_j P_j') \\ 2\delta_1 P_1 P_1' + 2\sigma_g^2 A P_1 P_1' + 2\sigma_g^2 S P_1 P_1' + 2\sigma_e^2 P_1 P_1' \\ \vdots \\ 2\delta_k P_k P_k' + 2\sigma_g^2 A P_k P_k' + 2\sigma_g^2 S P_k P_k' + 2\sigma_e^2 P_k P_k' \end{bmatrix}$$

$$= 2 \begin{bmatrix} trA^2 & trAS & trA & P_1AP_1' & \dots & P_kAP_k' \\ trSA & trS^2 & trS & P_1SP_1' & \dots & P_kSP_k' \\ trA & trS & trI & P_1P_1' & \dots & P_kP_k' \\ P_1AP_1' & P_1SP_1' & P_1P_1' & P_1P_1' & \dots & 0 \\ \vdots & & & & & \\ P_kAP_k' & P_kSP_k' & P_kP_k' & 0 & \dots & P_kP_k' \end{bmatrix} \begin{bmatrix} \sigma_g^2 \\ \sigma_g^2 \\ \sigma_e^2 \\ \delta_1 \\ \vdots \\ \delta_k \end{bmatrix}$$

The zero condition becomes

$$\begin{bmatrix} trA^2 & trAS & trA & P_1AP_1' & \dots & P_kAP_k' \\ trSA & trS^2 & trS & P_1SP_1' & \dots & P_kSP_k' \\ trA & trS & trI & P_1P_1' & \dots & P_kP_k' \\ P_1AP_1' & P_1SP_1' & P_1P_1' & P_1P_1' & \dots & 0 \\ \vdots & & & & & \\ P_kAP_k' & P_kSP_k' & P_kP_k' & 0 & \dots & P_kP_k' \end{bmatrix} \begin{bmatrix} \sigma_g^2 \\ \sigma_g^2 \\ \sigma_e^2 \\ \delta_1 \\ \vdots \\ \delta_k \end{bmatrix} = \begin{bmatrix} yAy \\ ySy \\ yy \\ yP_1P_1y \\ \vdots \\ yP_kP_ky \end{bmatrix}$$

Moving the $\delta_j$ across the equation

$$\begin{bmatrix} trA^2 & trAS & trA \\ trSA & trS^2 & trS \\ trA & trS & trI \\ P_1AP_1' & P_1SP_1' & P_1P_1' \\ \vdots & & \\ P_kAP_k' & P_kSP_k' & P_kP_k' \end{bmatrix} \begin{bmatrix} \sigma_g^2 \\ \sigma_g^2 \\ \sigma_e^2 \end{bmatrix} = \begin{bmatrix} yAy - \sum \delta_j P_j AP_j' \\ ySy - \sum \delta_j P_j SP_j' \\ yy - \sum \delta_j P_j P_j' \\ yP_1P_1y - \delta_1 P_1 P_1' \\ \vdots \\ yP_kP_ky - \delta_k P_k P_k' \end{bmatrix}$$

Plugging in

$$\delta_j P_j P_j = yP_jP_jy - \sigma_g^2 P_j AP_j' - \sigma_g^2 P_j SP_j - \sigma_e^2 P_j P_j'$$

we get

$$\begin{bmatrix} trA^2 & trAS & trA \\ trSA & trS^2 & trS \\ trA & trS & trI \\ P_1AP_1' & P_1SP_1' & P_1P_1' \\ \vdots & & \\ P_kAP_k' & P_kSP_k' & P_kP_k' \end{bmatrix} \begin{bmatrix} \sigma_g^2 \\ \sigma_g^2 \\ \sigma_e^2 \end{bmatrix} = \begin{bmatrix} yAy - \sum \delta_j P_j AP_j' \\ ySy - \sum \delta_j P_j SP_j' \\ yy - \sum \delta_j P_j P_j' \\ yP_1P_1y - (yP_1P_1y - \sigma_g^2 P_1AP_1' - \sigma_g^2 P_1SP_1 - \sigma_e^2 P_1P_1') \\ \vdots \\ yP_kP_ky - (yP_kP_ky - \sigma_g^2 P_kAP_k' - \sigma_g^2 P_kSP_k - \sigma_e^2 P_kP_k') \end{bmatrix}$$

The lower portion of the matrix is a tautolgy allowing the reduction to

$$\begin{bmatrix} trA^2 & trAS & trA \\ trSA & trS^2 & trS \\ trA & trS & trI \end{bmatrix} \begin{bmatrix} \sigma_g^2 \\ \sigma_g^2 \\ \sigma_e^2 \end{bmatrix} = \begin{bmatrix} yAy - \sum \delta_j P_j A P_j' \\ ySy - \sum \delta_j P_j S P_j' \\ yy - \sum \delta_j P_j P_j' \end{bmatrix}$$

$$\text{Let, } s_j := P_j S P_j' \ \ t_j := P_j A P_j' \ \ u_j := y P_j P_j' y$$

Note, this makes

$$\delta_j = u_j - \sigma_g^2 t_j - \sigma_g^2 s_j - \sigma_e^2$$

Collecting variance terms
Simplifying some of the traces

$$\begin{bmatrix} \sigma_g^2 \\ \sigma_g^2 \\ \sigma_e^2 \end{bmatrix} = \begin{bmatrix} trA^2 - \sum t_j^2 & trAS - \sum s_j t_j & trA - \sum t_j \\ trSA - \sum t_j s_j & \sum n_i^2 - \sum s_j^2 & N - \sum s_j \\ trA - \sum t_j & N - \sum s_j & N - k \end{bmatrix}^{-1} \begin{bmatrix} yAy - \sum u_j t_j \\ ySy - \sum u_j s_j \\ yy - \sum u_j \end{bmatrix}$$

#### A.1 Consistency of new estimator

The new estimator uses method of moments estimator and so therefore falls under M-estimation or generalized estimating equations theory. Assuming the mean only depends on the demographic variables used we know that $E(Y_i - X_i\beta)X_i = 0$ the model for the first moment is properly defined. For the second moment, the estimating equation $\psi$ is

$$\psi(Z_{ii'}, X_{ii'}, \beta) = (Z_{ii'} - X_{ii'}'\beta)X_{ii'}$$

Letting $Z = YY'$ and letting $X_{ii'}$ be the vector of the ijth element from $A$, $S$, and $I$. Assuming the site effects can be modeled as random mean, the expectation is

$$\begin{aligned} E(Z_{ii'} - X_{ii'}\beta)X_{ii'} &= E(E(Z_{ii'} - X_{ii'}\beta_0)X_{ii'}|X_{ii'}) \\ &= E(X_{ii'}\beta_0 - X_{ii'}\beta_0)X_{ii'}|X_{ii'} \\ &= 0 \end{aligned}$$

Making this a consistent estimator.

#### A.2 Consistency of other methods

For most of these methods, considering the first moment is sufficient to prove lack of consistency in general.

Removing the site effect by itself.

$$E(Y_i - X_{si}\beta_s)X_{si} = (X_d\beta_d - X_{si}\beta_s)X_{si}$$
$$X_d\beta_d = X_{si}\beta_s$$

which is not in general true for all i meaning this will not be a consistent estimator unless both effects on the mean are 0 or there is perfect separation of genetic ancestries between sites. This encompasses SWD, COMBAT, and COVBAT methods where the latter two methods approximate the site effect $\beta_s$ with a bayesian estimate.

Removing the site and demographic effects jointly

$$\begin{aligned} E(Y_i - X_{sdi}\beta_{sd})X_{sdi} &= (X_d\beta_d - X_{sdi}\beta_{sd})X_{sdi} \\ X_{di}\beta_d &= X_{si}\beta_s + X_{di}\beta_d \\ Z_i &= Y_i - X_{di}\beta_d - X_{si}\beta_s \\ E(Z_iZ_j - V_{ii'}\sigma)V_{ii'} &= 0 \\ A_{ii'}\sigma_g^2 + S_{ii'}\sigma_s^{2(R)} + I_{ii'}\sigma_e^2 &= A_{ii'}\sigma_g^2 + I_{ii'}\sigma_e^2 \end{aligned}$$

So the characterizing equation for the first moment yields an expectation of zero since the site effect is expected to be zero with the site effects being random effects. However, in the second moment, the residual site effects (that results from the collinearity with the genetic clusters, $\sigma_s^{2(R)}$ would lead to an imbalance in the characterizing equation meaning it would to be equivalent to 0 for all i, meaning this estiamtor, wouldn't guarantee consistency.

#### A.3 Variance and Wald test

First to include the uncertainty associated with estimating $\sigma_S^2$, we express the computation of heritability as $h^2 = f(\sigma_p^2, \sigma_g^2, \sigma_S^2) = \frac{\sigma_g^2}{\sigma_p^2 - \sigma_S^2}$, where $\sigma_p^2$ is the total variance of the phenotype ($\sigma_p^2 = \sigma_g^2 + \sigma_S^2 + \sigma_e^2$). Extending results from Ge et al [43] to multiple sites the variance of the random effects estimator is given by applying delta method

$$\begin{aligned} var(\hat{h}^2) = var(f(\sigma)) &\approx \frac{df(\sigma)}{d\sigma} var(\sigma) \frac{df(\sigma)}{d\sigma'} \\ \frac{df(\sigma)}{d\sigma} &= (\sigma_p^2 - \sigma_S^2)^{-2}(-\sigma_g^2, \sigma_p^2 - \sigma_S^2, \sigma_g^2) \end{aligned}$$

Without subpopulations

$$\begin{bmatrix} \sigma_g^2 \\ \sigma_s^2 \\ \sigma_e^2 \end{bmatrix} = \begin{bmatrix} trA^2 & trAS & trA \\ trSA & \sum n_i^2 & N \\ trA & N & N \end{bmatrix}^{-1} \begin{bmatrix} yAy \\ ySy \\ yy \end{bmatrix}$$

Let D be the determinant

$$\begin{bmatrix}\sigma_g^2\\ \sigma_s^2\\ \sigma_e^2\end{bmatrix} = 1/D \begin{bmatrix} N(\sum n_i^2 - 1)) & N(trA - trAS) & NtrAS - trA(\sum n_i^2) \\ N(trA - trAS) & NtrA^2 - tr^2A & trAStrA - NtrA^2 \\ NtrAS - trA(\sum n_i^2) & trAStrA - NtrA^2 & trA^2(\sum n_i^2) - tr^2AS \end{bmatrix} \begin{bmatrix} yAy \\ ySy \\ yy \end{bmatrix}$$

$$= \frac{1}{D} \begin{bmatrix} y'(N(\sum n_i^2 - 1)A + N(trA - trAS)S + NtrAS - trA(\sum n_i^2))y \\ y'(N(trA - trAS)A + (NtrA^2 - tr^2A)S + trAStrA - NtrA^2)y \\ y'((NtrAS - trA(\sum n_i^2))A + (trAStrA - NtrA^2)S + trA^2(\sum n_i^2) - tr^2AS)y \end{bmatrix}$$
$$\approx \frac{1}{D} \begin{bmatrix} y'(N(\sum n_i^2)A + N(trA - trAS)S + NtrAS - trA(\sum n_i^2))y \\ y'(N(trA - trAS)I + (\kappa - \tau)S + trAStrA - NtrA^2)y \\ y'((NtrAS - trA(\sum n_i^2))I + (trAStrA - NtrA^2)S + trA^2(\sum n_i^2) - tr^2AS)y \end{bmatrix}$$
$$\text{(using } \textstyle\sum n_i^2 - 1 \approx \sum n_i^2 \text{ and } trA^2 - tr^2A/N = N(\kappa - \tau))$$
$$\approx \frac{N^2}{D} \begin{bmatrix} y'Q_Ay \\ y'Q_Sy \\ y'Q_Ey \end{bmatrix}$$
$$= \frac{1}{D} \begin{bmatrix} y'Q_Ay \\ y'Q_Sy \\ y'Q_Ey \end{bmatrix}$$

$$var(\hat{h}^2) =\approx \frac{2}{(\sigma_p^2 - \sigma_s^2)^2}(-\sigma_g^2, \sigma_p^2 - \sigma_S^2, \sigma_g^2) \begin{bmatrix} 1 & 1 & 1 \\ 1 & 1 & -1 \\ 1 & -1 & 1 \end{bmatrix} (-\sigma_g^2, \sigma_p^2$$
$$= \frac{2}{(\sigma_p^2 - \sigma_S^2)^2} \left[(-\sigma_g^2 + \sigma_p^2 - \sigma_S^2 + \sigma_g^2, -\sigma_g^2 + \sigma_p^2 - \sigma_S^2 - \sigma_g^2, -\sigma_g^2 - \sigma_p^2 + \sigma_s^2 + \sigma_g^2)\right] (-\sigma_g^2, \sigma_p^2$$
$$= \frac{2}{(\sigma_p^2 - \sigma_S^2)^2} \left[(\sigma_p^2 - \sigma_S^2, -2\sigma_g^2 + \sigma_p^2 - \sigma_S^2, -\right.$$

$$Var(\sigma_p, \sigma_g, \sigma_e = \begin{bmatrix} Var(\sigma_p) & Cov(\sigma_p, \sigma_g) & \end{bmatrix}$$

$$\frac{2}{(\sigma_p^2 - \sigma_S^2)^2}(-2\sigma_p^2\sigma_g^2 + \sigma_S^2\sigma_g^2 + (\sigma_p^2 - \sigma_S^2)(-2\sigma_g^2 + \sigma_p^2 - \sigma_S^2))$$
$$\frac{2}{(\sigma_p^2 - \sigma_S^2)^2}(-4\sigma_p^2\sigma_g^2 + (\sigma_p^2 - \sigma_S^2)(\sigma_p^2 - \sigma_S^2))$$

$$\frac{2}{(\sigma_p^2-\sigma_S^2)^2}(-4\sigma_p^2\sigma_g^2+\sigma_p^4-2\sigma_p^2\sigma_S^2+\sigma_S^4)$$

The approximation assumes $trAS = trA/N_s, \sum n_i^2 - 1 = \sum n_i^2, \sum n_i^2 = N_s\bar{N}^2$ and same as Ge et al, $A \approx I$. Let $N(\sum n_i^2 - 1) := a, b = N(trA - trAS), c = NtrAS - trA(\sum n_i^2), d = N(trA - trAS), e = NtrA^2 - tr^2A), f = trAStrA - NtrA^2, g = NtrAS - trA(\sum n_i^2), h = trAStrA - NtrA^2, i = trA^2(\sum n_i^2) - tr^2AS$.

$$cov(\hat{\sigma}) = 2\begin{bmatrix} tr(Q_A Var(Y)Q_A Var(Y) & tr(Q_s Var(Y)Q_S Var(Y) & tr(Q_E Var(Y)Q_E Var(\text{[illegible]} \\ tr(Q_S Var(Y)Q_A Var(Y) & tr(Q_S Var(Y)Q_S Var(Y)) & tr(Q_S Var(Y)Q_E Var(\text{[illegible]} \\ tr(Q_E Var(Y)Q_A Var(Y)) & tr(Q_E Var(Y)Q_S Var(Y)) & tr(Q_E Var(Y)Q_E Var(\text{[illegible]} \end{bmatrix}$$

Assuming that $Var(Y) = \sigma_g^2 A + \sigma_S^2 S + \sigma_e^2 I \approx \sigma_p^2 I$.

$$cov = 2\sigma_p^2 \begin{bmatrix} tr(Q_A^2) & tr(Q_A Q_S) & tr(Q_A Q_E) \\ tr(Q_S Q_A) & tr(Q_S^2) & tr(Q_S Q_E) \\ tr(Q_E Q_A) & tr(Q_E Q_S) & tr(Q_E^2) \end{bmatrix}$$

$$\frac{2\sigma_p^2}{\nu_k}\begin{bmatrix} 1 & ? & -1 \\ ? & ? & ? \\ -1? & 1 & \end{bmatrix}$$

$$var(\sigma_g^2, \sigma_S^2, \sigma_e^2) = 2\sum \begin{bmatrix} tr(AVar(Y)AVar(Y)) & tr(AVar(Y)Var(Y)) & tr(AVar(Y)IVar(\text{[illegible]} \\ tr(Var(Y)AVar(Y)) & tr(Var(Y)Var(Y)) & tr(Var(Y)IVar(\text{[illegible]} \\ tr(AVar(Y)IVar(Y)) & tr(IVar(Y)Var(Y)) & tr(IVar(Y)IVar(\text{[illegible]} \end{bmatrix}$$

(Following simplification from Ge for symmetric matrices)

$$= 2tr(\sigma_P^2)\begin{bmatrix} tr(A^2) & tr(A) \\ tr(A) & N \end{bmatrix} \qquad (A \approx I)$$

$$\hat{v}ar(\hat{h}^2) = \frac{2tr(\sigma_P^2)}{(\sigma_P^2)^4}(\sigma_e^2, -\sigma_g^2)\begin{bmatrix} tr(A^2) & tr(A) \\ tr(A) & N \end{bmatrix}\begin{bmatrix} \sigma_e^2 \\ -\sigma_g^2 \end{bmatrix}$$
$$= \frac{2tr(\sigma_P^2)}{(\sigma_P^2)^4}(\sigma_e^2, -\sigma_g^2)\begin{bmatrix} tr(A^2) & tr(A) \\ tr(A) & N \end{bmatrix}\begin{bmatrix} \sigma_e^2 \\ -\sigma_g^2 \end{bmatrix}$$

Which is identical to the fixed effects model as shown in [43]. This means the variance is given by $var(\hat{h}^2) \approx 2/(trA^2 - tr^2A))$ and the wald test is given by

| | | Race | | | | |
|---|---|---|---|---|---|---|
| Site | Female | Asian | Black | Hispanic | White | Other |
| 1 | 111 (45%) | 8 (3.2%) | 16 (6.4%) | 148 (59%) | 47 (19%) | 30 (12%) |
| 2 | 38 (45%) | 2 (2.4%) | 0 (0%) | 13 (15%) | 65 (77%) | 4 (4.8%) |
| 3 | 213 (46%) | 0 (0%) | 64 (14%) | 348 (74%) | 39 (8.3%) | 17 (3.6%) |
| 4 | 234 (46%) | 3 (0.6%) | 67 (13%) | 103 (20%) | 222 (43%) | 117 (23%) |
| 5 | 121 (47%) | 1 (0.4%) | 67 (26%) | 9 (3.5%) | 152 (60%) | 26 (10%) |
| 6 | 227 (51%) | 8 (1.8%) | 15 (3.3%) | 75 (17%) | 296 (66%) | 55 (12%) |
| 7 | 114 (47%) | 6 (2.4%) | 44 (18%) | 36 (15%) | 137 (56%) | 22 (9.0%) |
| 8 | 106 (45%) | 32 (13%) | 3 (1.3%) | 54 (23%) | 104 (44%) | 45 (19%) |
| 9 | 151 (44%) | 12 (3.5%) | 32 (9.4%) | 129 (38%) | 116 (34%) | 51 (15%) |
| 10 | 225 (44%) | 11 (2.2%) | 20 (3.9%) | 306 (60%) | 137 (27%) | 34 (6.7%) |
| 11 | 160 (49%) | 2 (0.6%) | 63 (19%) | 32 (9.8%) | 194 (60%) | 35 (11%) |
| 12 | 195 (45%) | 10 (2.3%) | 151 (35%) | 37 (8.6%) | 176 (41%) | 55 (13%) |
| 13 | 257 (48%) | 7 (1.3%) | 97 (18%) | 45 (8.4%) | 321 (60%) | 65 (12%) |
| 14 | 54 (47%) | 1 (0.9%) | 2 (1.7%) | 6 (5.2%) | 95 (82%) | 12 (10%) |
| 15 | 170 (48%) | 2 (0.6%) | 192 (54%) | 14 (3.9%) | 94 (26%) | 53 (15%) |
| 16 | 269 (43%) | 1 (0.2%) | 7 (1.1%) | 74 (12%) | 504 (81%) | 36 (5.8%) |
| 17 | 216 (48%) | 9 (2.0%) | 3 (0.7%) | 17 (3.8%) | 396 (87%) | 28 (6.2%) |
| 18 | 142 (45%) | 6 (1.9%) | 29 (9.3%) | 26 (8.3%) | 219 (70%) | 33 (11%) |
| 19 | 47 (53%) | 0 (0%) | 31 (35%) | 5 (5.6%) | 42 (47%) | 11 (12%) |
| 20 | 96 (45%) | 0 (0%) | 95 (44%) | 5 (2.3%) | 91 (43%) | 23 (11%) |
| 21 | 181 (45%) | 5 (1.2%) | 63 (16%) | 99 (25%) | 206 (51%) | 28 (7.0%) |
| 22 | 20 (61%) | 2 (6.1%) | 5 (15%) | 6 (18%) | 16 (48%) | 4 (12%) |

**Table B1** Characteristics of ABCD dataset across sites

$$\frac{\hat{h}^2_{SNP}}{var(\hat{h}^2_{SNP}} \sim 1\chi^2_0 + 1/2\chi^2_1$$

### Appendix B ABCD characterization

### Appendix C Equivalence between observed GRM and true GRM

Due to differences in allelic frequency between different genetic clusters the estimated GRM ($A$) contains information on genetic ancestry in addition to the true GRM ($A^*$). In order to account for this we first consider the composition of the standardized genome $Z$ vs the true standardized genome $Z^*$. For the ith indivdiaul from the kth site and SNP

$$z^*_{kis} = \frac{x_{is} - 2p_{ks}}{f_s} \qquad \text{(True)}$$

$$z_{is} = \frac{x_{ii'} - 2p_s}{f_s}$$

Letting $a_{ks} := \frac{2(p_{ks}-p_s}{f_s}$ and $p_s = 1/N(n_1p_{1s}+n_2p_{2s})$ we can rearrange this as

$$z_{is} = z^*_{kis} + a_{ks}$$

Therefore the true and estimated GRMs can be related as

$$A^* = A - \frac{1}{M}(aZ' + Za' - aa')$$

For the case of two clusters where they can be related by 1 PC we have the projection

$$Q_{PC} = I - \begin{bmatrix} v_1^2\mathbf{1_{n_1,n_1}} & -v_1v_2\mathbf{1}_{n_1,n_2} \\ -v_1v_2\mathbf{1}_{n_2,n_1} & v_2^2\mathbf{1}_{n_2,n_2} \end{bmatrix}$$
$$\text{Where } v_k = \frac{1}{n_k}\sqrt{\frac{n_1n_2}{N}}$$

Apply this to the true GRM

$$Q_{PC}A^*Q_{PC} = Q_{PC}(A - \frac{1}{M}(aZ' + Za' - aa'))Q_{PC}$$

But since

$$Q_{PC}a = a - \begin{bmatrix} v_1^2\mathbf{1_{n_1,n_1}} & -v_1v_2\mathbf{1}_{n_1,n_2} \\ -v_1v_2\mathbf{1}_{n_2,n_1} & v_2^2\mathbf{1}_{n_2,n_2} \end{bmatrix} \begin{bmatrix} a_{1s}e_{n_1} \\ a_{2s}e_{n_2} \end{bmatrix} = 0$$

Therefore $Q_{PC}A^*Q_{PC} = Q_{PC}AQ_{PC}$

### Appendix D Variance of AdjHE Random site effects estimator

The variance of the random effects estimator is given by applying delta method

$$var(\hat{h}^2) = var(f(\sigma)) \approx \frac{df(\sigma)}{d\sigma}var(\sigma)\frac{df(\sigma)}{d\sigma'}$$
$$\frac{df(\sigma)}{d\sigma} = \frac{1}{(\sigma_P^2)^2}(\sigma_e^2, 0, -\sigma_g^2)$$

Noting that $df(\sigma)/d\sigma$ has a 0 as the second element, the 2nd row and 2nd column are inconsequential and we get

$$var(\hat{h}^2) = var(f(\sigma_g^2, \sigma_e^2)) \approx \frac{df(\sigma_g^2, \sigma_e^2)}{d\sigma} var(\sigma_g^2, \sigma_e^2) \frac{df(\sigma_g^2, \sigma_e^2)}{d\sigma'}$$

$$var(\sigma_g^2, \sigma_e^2) = 2 \sum \begin{bmatrix} tr(AVar(Y)AVar(Y)) & tr(AVar(Y)IVar(Y)) \\ tr(AVar(Y)IVar(Y)) & tr(IVar(Y)IVar(Y)) \end{bmatrix}$$

(Following simplification from Ge for symmetric matrices)

$$= 2tr(\sigma_P^2) \begin{bmatrix} tr(A^2) & tr(A) \\ tr(A) & N \end{bmatrix} \qquad (A \approx I)$$

$$var(h^2) = \frac{2}{\sigma_P^2}(\sigma_e^2, -\sigma_g^2) \begin{bmatrix} tr(A^2) & tr(A) \\ tr(A) & N \end{bmatrix} \begin{bmatrix} \sigma_e^2 \\ -\sigma_g^2) \end{bmatrix}$$

$$= \frac{2}{\sigma_P^2}(\sigma_e^2 trA^2 - \sigma_g^2 trA, \sigma_e^2 trA - \sigma_g^2 N) \begin{bmatrix} \sigma_e^2 \\ -\sigma_g^2 \end{bmatrix}$$

$$= 2\frac{(\sigma_e^2)^2 trA^2 - 2\sigma_g^2\sigma_e^2 trA + (\sigma_g^2)^2 N}{\sigma_g^2 + \sigma_e^2}$$

Which is identical to the fixed effects model. The wald test is then given by

$$\frac{\hat{h}^2_{SNP}}{var(\hat{h}^2_{SNP}} \sim 1\chi_0^2 + 1/2\chi_1^2$$

Extending results from https://www.nature.com/articles/ncomms13291 (Methods - Sampling variance of the point estimator) we see the variance of the fixed effects estimator is given by

$$var(\hat{h}^2_{SNP}) \approx \frac{2}{tr(A^2) - 2trA + n - \sum(PAP - 1)^2}$$

### Appendix E Biasedness not accounting for site

Consider a multisite study in which the dummy matrix codifying at which site a given measurement was taken ($X_S$). In the AdjHE model these would be accounted for as a fixed effect. However, it is supposed that the site matrix breaks the assumption of the AdjHE model, speciiically that $X_S \not\perp X_G$. The variance estimates given without considering site in the case that site does impact the phenotype is given by the AdjHE estimator as

$$\hat{\beta} = \left( \begin{bmatrix} A \\ I \end{bmatrix} [A \; I] \right)^{-1} \begin{bmatrix} A \\ I \end{bmatrix} YY'$$

Then the expectation of our estimator is

$$\begin{aligned} E\hat{\sigma} &= \left( \begin{bmatrix} A \\ I \end{bmatrix} [A \; I] \right)^{-1} \begin{bmatrix} A \\ I \end{bmatrix} EYY' \\ &= \left( \begin{bmatrix} A \\ I \end{bmatrix} [A \; I] \right)^{-1} \left( \left( \begin{bmatrix} A \\ I \end{bmatrix} [A \; I] \right) \begin{bmatrix} \sigma_g^2 \\ \sigma_e^2 \end{bmatrix} + \begin{bmatrix} A \\ I \end{bmatrix} S\sigma_g^2 \right) \\ &= \begin{bmatrix} \sigma_g^2 \\ \sigma_e^2 \end{bmatrix} + \left( \begin{bmatrix} A \\ I \end{bmatrix} [A \; I] \right)^{-1} \begin{bmatrix} trAS \\ n \end{bmatrix} \sigma_g^2 \\ &= \begin{bmatrix} \sigma_g^2 \\ \sigma_e^2 \end{bmatrix} + \frac{1}{ntrA^2 - (trA)^2} \begin{bmatrix} ntrAS - ntrA \\ ntrA^2 - trAStrA \end{bmatrix} \sigma_g^2 \end{aligned}$$

This is unbiased in two casees:

1. When $\sigma_g^2 = 0$
2. When
$$\begin{bmatrix} ntrAS - ntrA \\ ntrA^2 - trAStrA \end{bmatrix} = 0$$

For the second case this is when

$$trAS = trA \text{ and } ntrA^2 = trAStrA$$

Which is the same as

$$ntrA^2 = (trA)^2$$

Thanks to consultation with Robert Lewis on StackExchange post https://math.stackexchange.com/questions/506962/expressing-the-trace-of-a2-by-trace-of-a we know this is true when

$$C(A) = (trA)^2 \frac{n-1}{2n}$$

Where $C(A)$ is the sum of the product of all unique eigenvalue pairs of A. When $trAS > trA, \sigma_g^2$ will be overestimated and when $ntrA^2 > trAStrA, \sigma_e^2$ will be overestimated.

### Appendix F Simulation Results

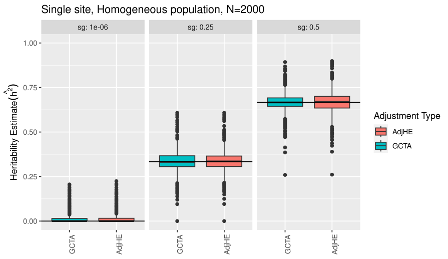


**Fig. F1** Heritability estimates based on 100 simulations repeated across an array of simulated heritabilities (black line) for sample sizes of 2000 for a homogeneous population at a single site.

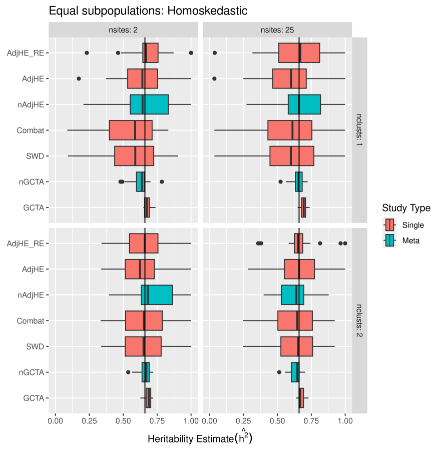


**Fig. F2** Simulated phenotypic variance with an increasing number of sites (columns) and clusters (rows). The total phenotypic variance was simulated 50% from genetic data, 25% from sites, and 25% from noise making the simulated heritability 66% (vertical black line).

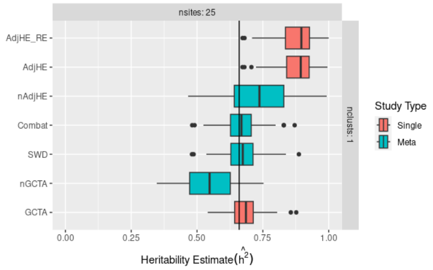


**Fig. F3** Simulated phenotypic variance with fixed site effects evenly spaced with mean zero. The total phenotypic variance was simulated 50% from genetic data, 25% from sites, and 25% from noise making the simulated heritability 66% (vertical black line).

### Appendix G ABCD Results

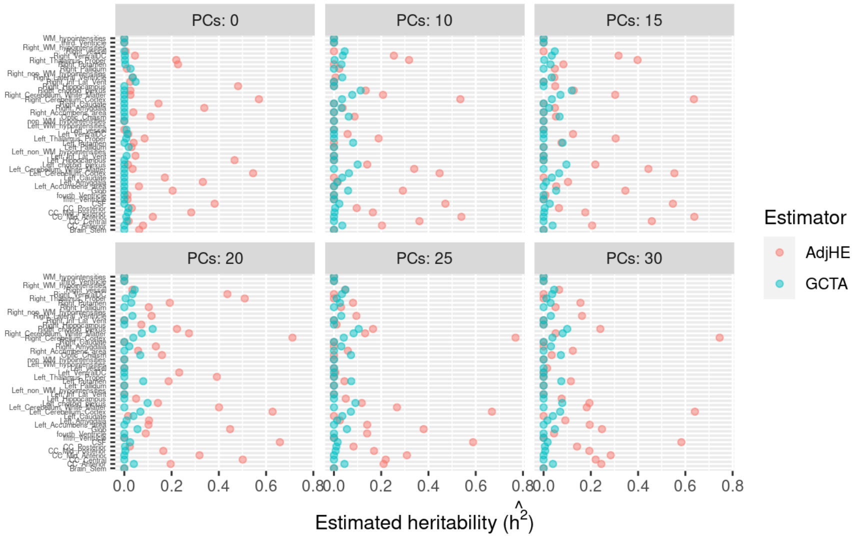


**Fig. G4** Heritability estimates for site 16 across all ancestries. Each IDP is represented along the y-axis. GCTA estimates are Blue whereas AdjHE estimates are red. The plots are faceted by the number of PCs controlled for 0, 5, 10, 15, 20, 25, 30.

### Appendix H PCA R2

##### H.0.1 Potential bias not accounting for sites

Not accounting for site effects can bias heritability estimates. Following from Section E, the expectation of our estimates without controlling for the site effect is

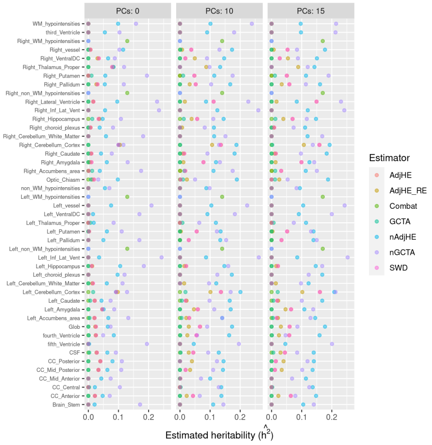


**Fig. G5** Heritability estimates for the full European ancestry from all sites dataset for multiple areas of the brain by all estimators. The columns are faceted by the number of PCs controlled for 0, 5, and 10.

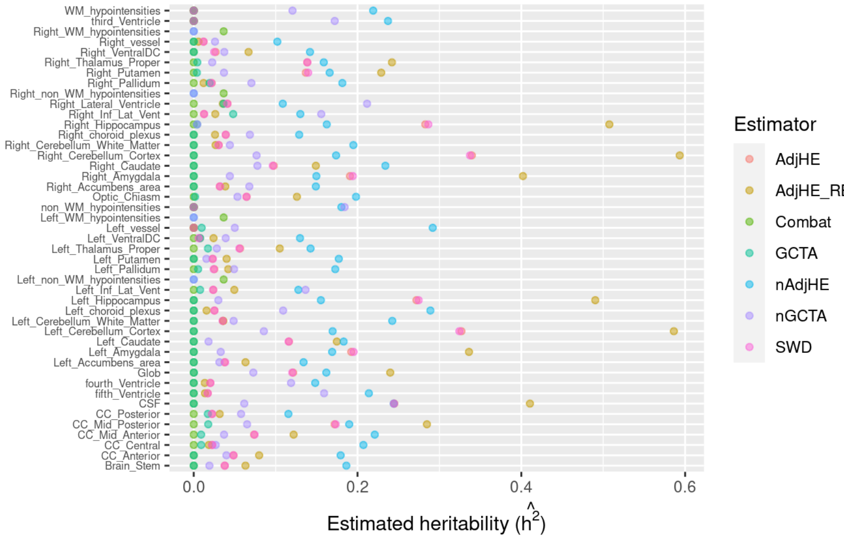


**Fig. G6** Heritability estimates for the full ABCD dataset for multiple areas of the brain (y-axis) by multiple estimators represented as different colors.

$$E\begin{bmatrix}\hat{\sigma}_g^2\\ \hat{\sigma}_e^2\end{bmatrix} = \begin{bmatrix}\sigma_g^2\\ \sigma_e^2\end{bmatrix} + \frac{1}{ntr\boldsymbol{A}^2-(tr\boldsymbol{A})^2}\begin{bmatrix}ntr\boldsymbol{AS}-ntr\boldsymbol{A}\\ ntr\boldsymbol{A}^2-tr\boldsymbol{AS}tr\boldsymbol{A}\end{bmatrix}\sigma_g^2$$

| Size | AdjHE | AdjHE (RE) | GCTA |
|---|---|---|---|
| 500 | 0.05 | 0.08 | 0.29 |
| 1000 | 0.08 | 0.33 | 1.42 |
| 2500 | 0.77 | 4.66 | 17 |
| 5000 | 5.6 | 35 | 109 |
| x faster than GCTA | $\approx 20$ | $\approx 3-4$ | 1 |

**Table G2** Average estimation time averaged over 100 simulations. The last row compares the average ratio of GCTA estimation over both AdjHE methods and reports the relative speed up.

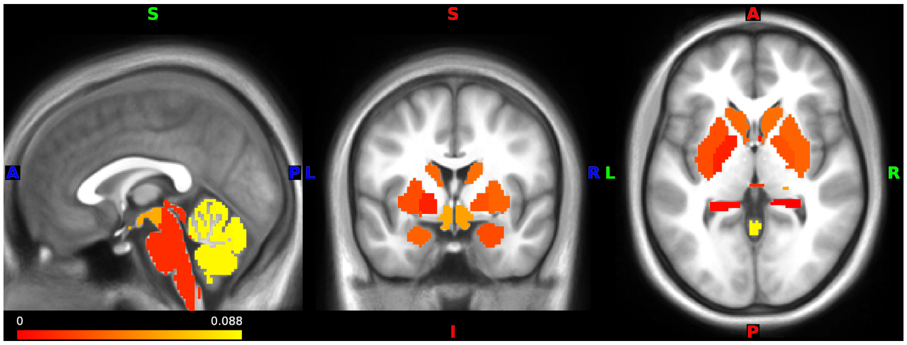


**Fig. H7** $R^2$ of the first 20 PCs projected onto the brain.

Since the sample size is implicit in the traces as well, consistency is harder to track. Accounting for sites is therefore necessary in general to avoid biased estimates except in special cases when the site effects are zero or if the GRMs are balanced between sites. See Section E for situations under which heritability will be over or underestimated.
